# Neural representations of predicted events: Evidence from time-resolved EEG decoding

**DOI:** 10.1101/2024.01.05.574347

**Authors:** Ai-Su Li, Jan Theeuwes, Dirk van Moorselaar

**Affiliations:** Institute Brain and Behavior Amsterdam, Department of Experimental and Applied Psychology, Vrije Universiteit Amsterdam, Amsterdam, the Netherlands; Department of Psychology, Soochow University, Suzhou, China; William James Center for Research, ISPA-Instituto Universitario, Lisbon, Portugal

**Author notes:** Correspondence should be addressed to: Ai-Su Li, Department of Psychology, Soochow University, WenJing Street 1, 215000 Suzhou, China.; Dirk van Moorselaar, Department of Experimental and Applied Psychology, Vrije Universiteit Amsterdam, Van der Boechorststraat 7-9, 1081 BT Amsterdam, The Netherlands. Conflict of Interest: The authors declare no competing financial interests.

## Abstract

Through statistical learning, humans are able to extract temporal regularities, using the past to predict the future. Evidence suggests that learning relational structures makes it possible to anticipate the imminent future; yet, the neural dynamics of predicting the future and its time-course remain elusive. To examine whether future representations are denoted in a temporally discounted fashion, we used the high-temporal-resolution of electroencephalography (EEG). Observers were exposed to a fixed sequence of events at four unique spatial positions within the display. Using multivariate pattern analyses trained on independent pattern estimators, we were able to decode the spatial position of dots within full sequences, and within randomly intermixed partial sequences wherein only a single dot was presented. Crucially, within these partial sequences, subsequent spatial positions could be reliably decoded at their expected moment in time. These findings highlight the dynamic weight changes within the assumed spatial priority map and mark the first implementation of EEG to decode predicted, yet critically omitted events.

**Impact statement:** Utilizing high-temporal-resolution EEG, the dynamic weight changes of assumed spatial priority map were visualized by decoding the spatial position of expected, yet omitted, events at their expected moment in time.

## Introduction

An essential aspect of our capacity to perceive and interact with the visual environment lies in our ability to discern recurrent patterns from past experiences and utilize them to forecast future occurrences, thereby guiding our actions. Critically, our ever-evolving surroundings, filled with abundant information, maintain structure and stability. For instance, through extensive practice in specific ball sports, we acquire the skill to predict the ball’s trajectory, enabling us to anticipate its future position and precisely intercept it when the need arises. At the behavioral level, the benefits of such sequence learning is demonstrated through the serial reaction time (SRT) task pioneered by Nissen and Bullemer (1987), wherein a single stimulus is sequentially presented at various locations (e.g., 4). The critical finding is that, even without any explicit awareness, observers can track the current position of the target stimulus (using spatially compatible keys) faster when the target follows a particular repetitive sequence compared to a random sequence. Subsequent experiments demonstrated that this learning could not be attributed to response-response or stimulus-response associations (see Schwarb & Schumacher, 2012 for a review) as learning persisted even when the motor component was removed (Remillard, 2003). Thus, unintentionally, human observers learn perceptual regularities, leading to more efficient processing of expected sensory events.

In more recent years, it has become clear that such perceptual learning is not restricted to sequentially presented individual shapes and associated responses, as observers can also learn spatial associations across trials during visual search (e.g., Boettcher et al., 2022; Li et al., 2022, 2023; Li & Theeuwes, 2020; Li et al., under review; Toh et al., 2021). For example, in Li and Theeuwes (2020), the target appeared with equal probability across all display locations, yet critically, two across-trial regularities regarding target locations were introduced so that the target presented at a specific location (e.g., left side of the display) was always followed by a target at the opposite display location. While most participants were unable to explicitly report these across-trial regularities, performance on predictable trials was nevertheless facilitated (i.e., faster RTs and higher accuracies) compared to unpredictable trials. This type of learning was classified as a form of statistical learning — a process by which observers extract statistical regularities in the environment from past experiences (see Frost et al., 2019; Theeuwes et al., 2022 for a review). When observers optimize their visual search performance according to the learned regularities that exist in the display, it is often claimed that through statistical learning the weights within an assumed spatial priority map change such that attention becomes biased towards locations where the target appears with a higher probability (e.g., Fecteau & Munoz, 2006; Ferrante et al., 2018; Theeuwes et al., 2022; Zelinsky & Bisley, 2015). Notably, in across-trial statistical learning the regularity is occurring across trials suggesting that local weight changes within the assumed priority map dynamically adapt on the basis of the trial-to-trial probabilities.

In this perspective the attentional priority landscape is not inherently a static representation constructed by previous selection episodes, but instead it operates as a dynamic system, wherein attentional priority undergoes continual adjustments based on anticipations triggered by the current event. This dynamic view on attentional priority, wherein the current state predicts the upcoming (future) states, is reminiscent of the successor representation framework (SR, Dayan, 1993). This framework postulates a predictive representation wherein the current state is represented in terms of its future (successor) states, in a temporally discounted fashion. While the SR framework was originally conceptualized in the context of hippocampal representations (e.g., Dayan, 1993; Fang et al., 2023; Stachenfeld et al., 2017), a recent functional magnetic resonance imaging (fMRI) study by Ekman et al. (2023) showed that after visual sequence learning, not only the hippocampus, but also V1 receptive fields became tuned to respond to both the current input, as well as expected future events. The fact that reactivations of future sequence locations were also observed in early visual cortex suggests that the successor representations are a more ubiquitous coding schema than previously assumed.

The evidence above highlights that the relational structure, for instance that event A is usually followed by event B, that is learned through exposure during past experiences is not passively represented, but instead is actively used to anticipate the imminent future. Consistent with this, previous research has repeatedly shown that prior expectations influence neural activity in the visual cortex (e.g., Fiser et al., 2016; Kok et al., 2013; Kok et al., 2012). Yet, in the context of learned spatial associations the time course of these effects remains largely unexplored, and it thus also remains unclear how and when the current representation shapes future representations in the context of spatial associations. As Ekman et al. (2023) employed fMRI as a way to measure brain activity, the temporal dynamics of the observed reactivations remain elusive. Therefore, in the current study, we relied on the high temporal resolution of electrophysiology (EEG) to investigate the temporal dynamics of anticipating future events. Previous research has demonstrated that the expected neural representation induced by probability cues on a trial-by-trial basis is already activated prior to actual stimulus onset (Kok et al., 2017). Here, we specifically set out to test whether stimulus presentation within a structured sequence also serves as a cue that triggers the retrieval of successor locations.

To test this, we adopted the sequence learning task of Ekman et al. (2023). Participants were exposed to an arbitrary spatiotemporal sequence of dots (see Figure 1) consisting of four items (A-B-C-D). Critically, after a training phase with repeated exposure to the sequence, we randomly intermixed full sequences with partial sequences, wherein only a single stimulus of the sequence appeared on screen at its expected moment in time (e.g., - B - -). Using time-resolved multivariate pattern analyses, we were able to test whether merely presenting one of the items in a learned sequence would trigger activation at the successor locations (e.g., C and D) and critically for the current study, if so, at what moment in time.

**Figure 1.**
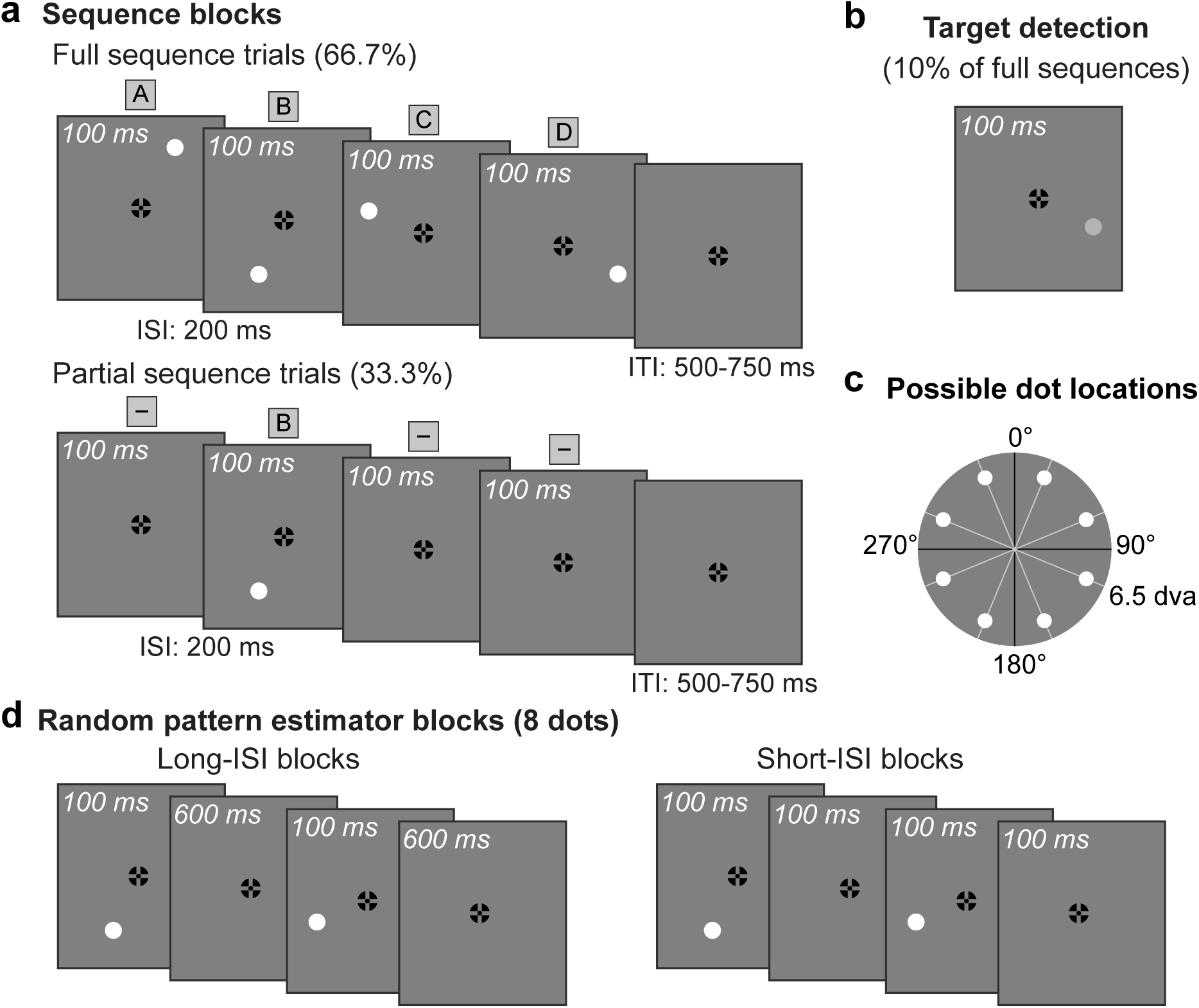
Stimuli and design in sequence blocks and pattern estimator blocks. (a) Illustration of sequence blocks containing full and partial sequence trials. In full sequence trials the white dot was presented for 100 ms at one of four positions successively in a fixed order with an inter-stimulus interval (ISI) of 200 ms. During partial sequence trials only one dot of the sequence (equal probability for each position in the sequence) was presented for 100 ms and all intervals remained the same as full sequence trials. (b) Illustration of target (i.e., gray dot detection) frame. (c) Graphical illustration of the eight possible dot locations, out of which only four were shown in full and partial sequence trials. (d) During random pattern estimator blocks, the white dot was presented at one of eight possible locations (as illustrated in Figure 1c) for 100 ms in random order with an ISI of 600 ms in long-ISI blocks and of 100 ms in short-ISI blocks.

## Results

While recording EEG and gaze behavior, participants (*N* = 25) had to detect each time a white dot turned grey within a fixed four-position dot sequence (i.e., ABCD; Figure 1a/b). Unbeknownst to participants these rare events only occurred at the final sequence position. Critically, after an initial exposure phase to familiarize participants with the sequence order, partial sequences, wherein only a single stimulus of the sequence was presented at its expected moment in time (e.g., -B--), were randomly intermixed into these full sequences (Figure 1a; see Methods section for details). This design allowed us to establish whether, and critically if so when, presenting a single item of a learned sequence would trigger the retrieval of successor locations within that sequence (Ekman et al., 2023).

### Within-block decoding: Increasing the interval between successive stimuli enhances performance

Before delving into the neural representations of specific positions within the sequence, including both present and absent dots, we initially ensured the robustness of decoding individual dot positions using data from independent pattern estimator blocks. Within these blocks participants were instructed to passively view a dot stimulus appearing at one of eight locations in random order (Figure 1c). Previous studies have successfully employed multivariate pattern analyses based on independent pattern estimators with either short (range: 0-200 ms; e.g., Alilovic et al., 2021; Blom et al., 2020; Robinson et al., 2020) or long (range: 433-1200 ms; e.g., Hogendoorn & Burkitt, 2018; Kok et al., 2017) ISI’s in between successive stimuli. Although these approaches have their own benefits (see Methods for details), here for the first time we attempted to decode the spatial position of expected dots in the absence of visual input. Given that this approach is arguably more challenging due to the event of interest not being time-locked to an evoked response, we chose to employ both types of pattern estimators to establish whether one type is more suited than the other. As visualized in Figure 2a, we observed that classifiers trained on data from both pattern estimators could decode individual dot positions well above chance level. However, noteworthy distinctions emerged between the two distinct pattern estimator conditions. The long-ISI pattern estimator decoding demonstrated superior overall classification performance, with subsequent peaks in decoding occurring at later time points (the short-ISI pattern estimator displayed an initial peak at approximately 88 ms, while the long-ISI pattern estimator showed an initial peak at around 135 ms). Previous research has established that multiple stimuli within a sequence can be simultaneously represented, with each individual stimulus being decodable for approximately one second (Carlson et al., 2013; King & Wyart, 2021). Arguably, the signal contamination stemming from preceding dots may also account for the observed differences between independent pattern estimators, with the sustained influence from preceding dots being more pronounced in the short-ISI pattern estimator blocks, where subsequent stimuli are temporally closer together compared to the long-ISI pattern estimator blocks. Critically, control analysis convincingly demonstrated that this above-chance classification could not be explained by systematic eye movements as the same analysis using the x, y coordinates measured by the eye tracker did not result in above-chance classification (e.g., Chota et al., 2023; Johnson et al., 2023).

**Figure 2.**
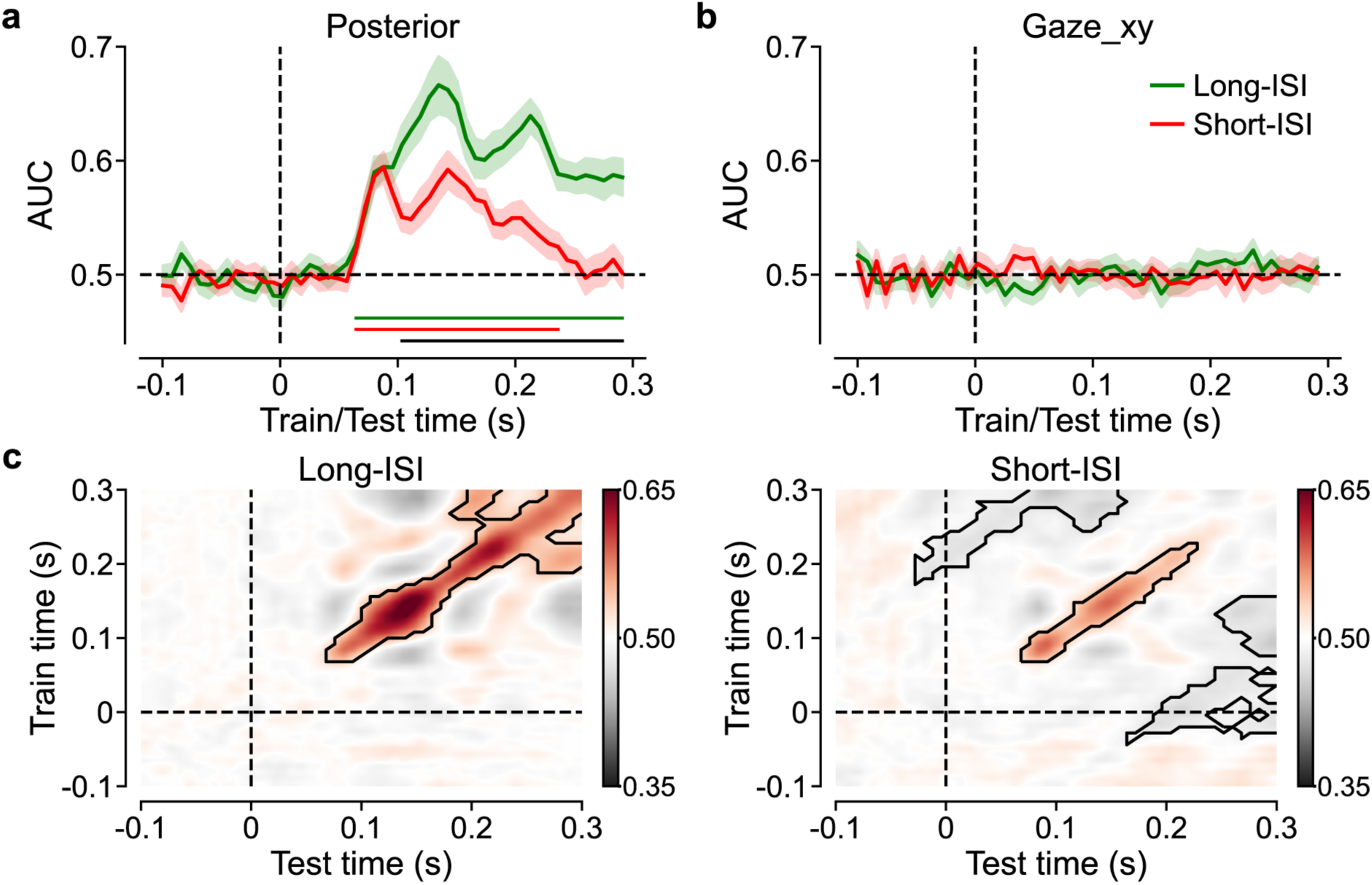
Pattern estimator position (four dots from the sequence) decoding results. The decoder was trained and tested at the same time point with posterior electrodes (a), or gaze positions (b) as features. Horizontal dashed lines indicate chance level while vertical dashed lines indicate dot onset. Green and red solid lines represent long- and short-ISI pattern estimators, respectively. Shaded region represents standard error of the mean (SEM, same applied to subsequent figures). Corresponding color bars below the x-axis indicate clusters that significantly differed from chance while the black bar indicates clusters with a significant difference between long- and short-ISI pattern estimators (*p* < .05). (c) Temporal generalization matrix, with posterior electrodes as features, showing classification performance as a function of training and test time point in long- and short-ISI blocks. Solid black lines indicate clusters that significantly different from chance (*p* < 0.05). All corresponding results that discriminated eight positions can be observed in Figure 2 – figure supplement 1.

In order to determine whether an individual event automatically triggers the retrieval of successive events and pinpoint the timing of this process, our study aimed to decode the spatial position of absent, yet anticipated dots. As noted, pattern estimator decoding displayed two successive peaks, and it was not immediately clear what cognitive processes these peaks represented, and which would be most applicable for position decoding within structured sequences. To gain a better understanding of the dynamics of position decoding, we therefore employed a generalization across time approach while applying the pattern classifiers (King & Dehaene, 2014), wherein a classifier trained at a specific time point is tested on all time points. The resulting generalization across time matrix (with training time on y-axis; testing time on x-axis) can help identify periods during which a representation remains stable, i.e., generalizes across time. As depicted in Figure 2c, both pattern estimators exhibited little to no evidence of early off-diagonal decoding, which has been associated with the perceptual maintenance of early sensory representations (e.g., Marti & Dehaene, 2017; Meijs et al., 2019; Weaver et al., 2019). Furthermore, in line with Hogendoorn and Burkitt (2018), the temporal generalization matrix revealed no above-chance classification performance at the intersections of the distinct peaks in diagonal decoding, suggesting that these peaks represent sequential, distinct patterns of neural activation (King & Dehaene, 2014).

Considering that our classifier likely tracked the dynamic transformation from representation to another over time, and we had no a priori prediction regarding the optimal moment to capture the processing of an anticipated, yet critically missing event, we decided not to restrain our cross-block classification (i.e., pattern estimator to structured sequence data classification) to a specific time window. Instead, we explored the entire time range using only the data from the long-ISI pattern estimator, as that one demonstrated superior decoding.

### Cross-block decoding: predecessor events in a sequence enhance decoding of the current event

Before examining the neural representation of anticipated, yet absent events, we first tested whether the data from the pattern estimator sequences could be used to decode present dot positions within the structured sequence. For this purpose, time courses of the pattern estimator blocks and the main task were aligned relative to dot onset and decoding was performed across time, with the model trained on pattern estimator data and tested on main task data, separate for full sequences (i.e., BCD) and partial sequences (e.g., -B--). Note that in this analysis, the first dot position (i.e., A), which does not allow to dissociate full from partial sequences, was not included. As shown in Figure 3a, cross-block classification also showed robust decoding of dot position, irrespective of whether the dot was embedded in a full sequence, or whether that dot was the only dot presented within the sequence (partial sequences). Notably, while clusters of significant decoding exhibited the same temporal dynamics, decoding was more pronounced approximately 200 ms following dot onset in the full sequence trials relative to partial sequence trials. Although speculative, we attribute this difference in decoding between two identical events to a more variable tuning of attention in the partial sequence relative to the full sequences. Whereas in the latter attentional shifts are yoked to a sequence of events that are fixed in time, attentional tuning arguably became more variable when dots were omitted from the sequence, and participants had to thus rely on internally generated representations. Consistent with this an exploratory analysis of the event-related waveforms elicited by posterior contralateral electrodes showed that especially the N1 component, which is linked to the orienting of attention to a task-relevant stimulus and modulated by the time in between successive events (Luck et al., 1990) was more pronounced in full than in partial sequences (Figure 3b). We then believe that this N1 modulation underlies our observed difference in decoding.

**Figure 3.**
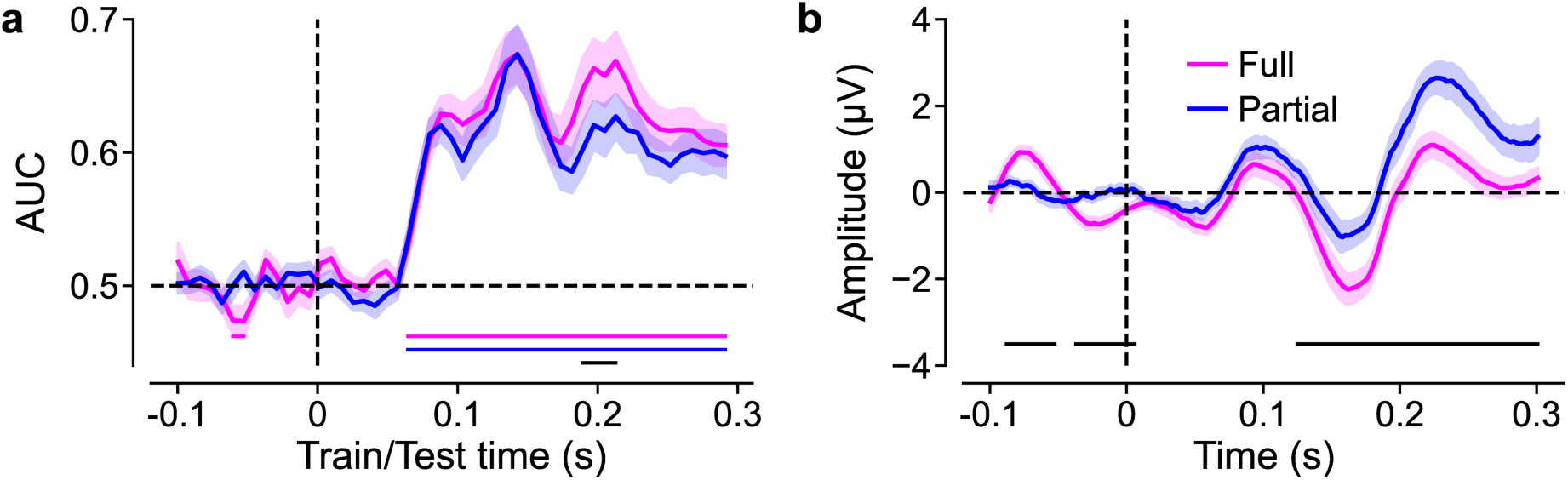
Position decoding and ERPs results for present dots. (a) Decoding of present dot positions (i.e., BCD) based on long-ISI pattern estimator (training and testing at the same time point) using posterior electrodes. (b) ERPs for the 8 posterior electrodes contralateral to each present dot position, collapsed over dots. The pink and blue lines represent full and partial sequence trials, respectively. Corresponding color bars below the x-axis indicate clusters that significantly differed from chance while black bars indicate clusters with a significant difference between full and partial sequence trials (*p* < .05).

### Cross-block decoding: Expected, but absent (immediate) successor events are represented at their expected moment in time

Having established that we could robustly decode dot positions within the structured sequences using independent pattern estimator data, we next examined whether those same spatial positions could be decoded when the anticipated dots were omitted from the sequence. For this purpose, following Ekman et al. (2023), we categorized omitted dots within partial sequences into two conditions: ‘predecessor’ and ‘successor’, based on their position in relation to the presented dot. For instance, in the partial sequence, -B--, A would be categorized as a predecessor position, while dots C and D would be designated as successor positions. As visualized in Figure 4, whereas decoding performance for predecessor locations was at chance level, despite the absence of evoked response classifiers were able to reliably classify successor locations, with above chance decoding emerging approximately around the time that the stimulus should have appeared. Counter to decoding of present dots however, which was largely restricted to the diagonal of temporal generalization matrix (Figure 4b), successor decoding was more diffuse (Figure 4c), arguably because the temporal predictions were not reset by actual stimulus presentation and hence somewhat variable. Next, we explored whether the observed successor decoding was observed independent of the temporal distance from the presented dot. As shown in Figure 5, whereas the successor representation immediately following the presented dot could be decoded reliably, there was little evidence that the subsequent successor representation was also evident in the ongoing EEG signal. Note that these results do not necessarily mean that only the immediate successor was actively represented, as EEG might simply not be sensitive enough to pick up on these weak and temporal diffuse representations. Indeed, the fMRI study by Ekman et al. (2023) showed that while all successor representations were represented, activations decreased with increasing temporal distance from the present dot. Nevertheless, the current results convincingly demonstrate that anticipated events are actively represented at their expected moments in time, even without an evoked response elicited by that event.

**Figure 4.**
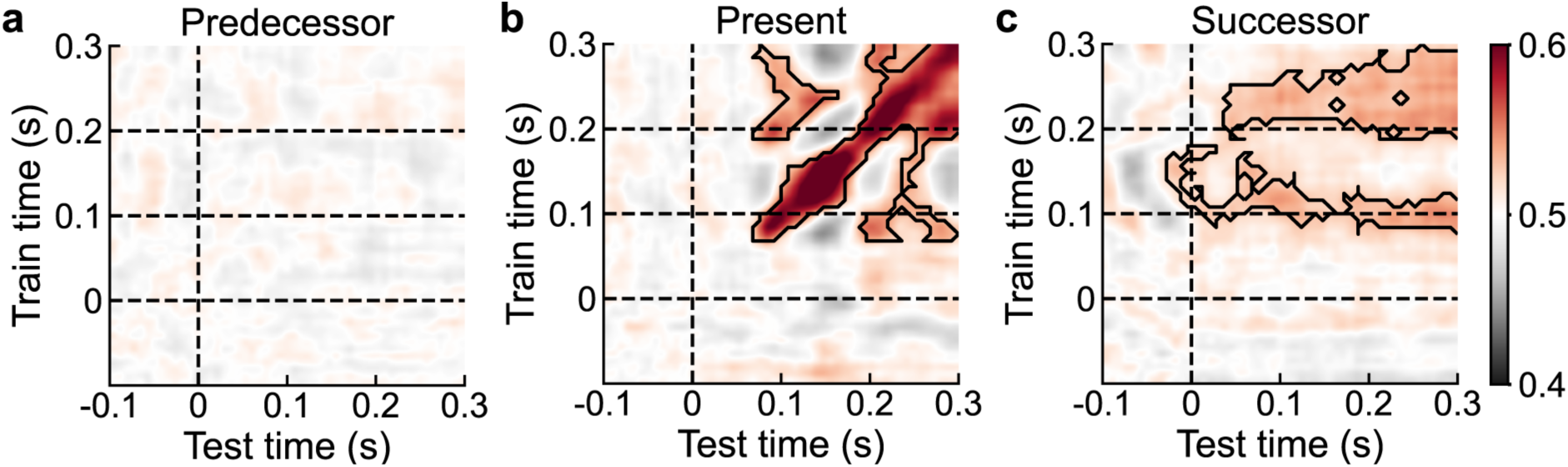
Position decoding results of partial sequence trials. Temporal generalization of decoding absent predecessor (a), present (b), and absent successor (c) positions of partial sequence trials based on long-ISI pattern estimator using posterior electrodes. Solid black lines indicate clusters that significantly different from chance (*p* < 0.05).

**Figure 5.**
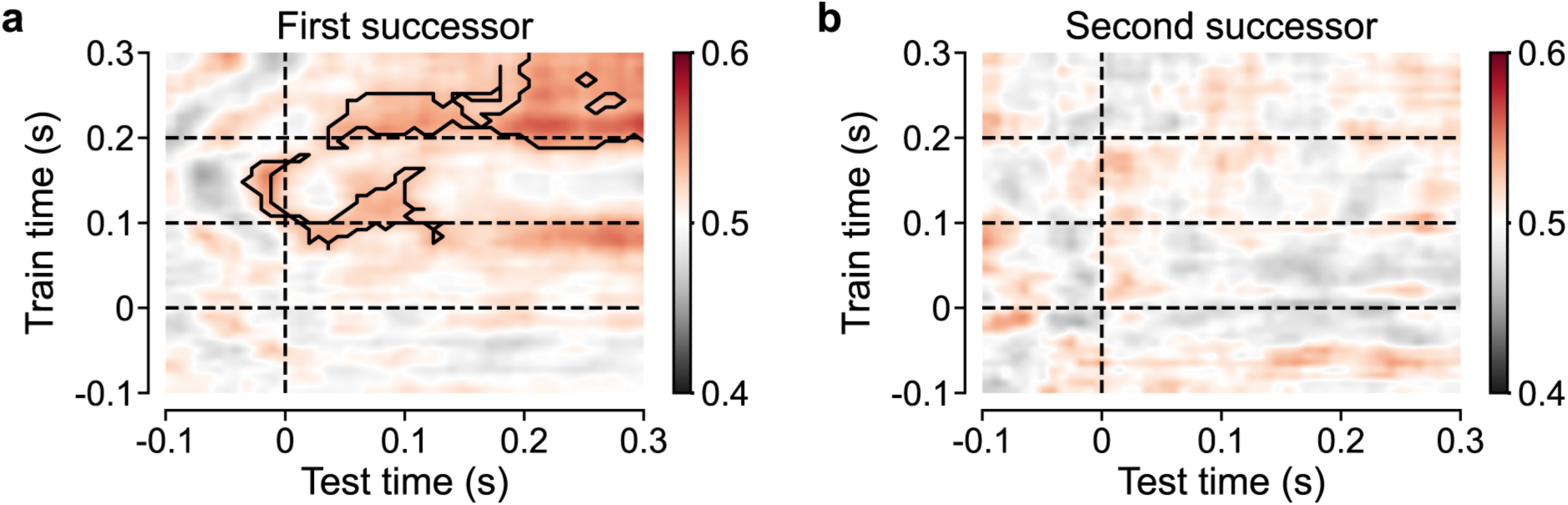
Successive position decoding results of partial sequence trials. Temporal generalization of decoding the first successor (a) and second successor (b) positions based on long-ISI pattern estimator using posterior electrodes. Solid black lines indicate clusters that significantly different from chance (*p* < 0.05).

## Discussion

Recently, a large body of research has shown that people are able to learn from past experiences and apply what is learned to optimize selection (e.g., Duncan et al., 2023; Ferrante et al., 2018; Gao & Theeuwes, 2020; Huang et al., 2022; Liesefeld & Müller, 2021; van Moorselaar & Theeuwes, 2023; Wang & Theeuwes, 2018; Xu et al., 2022). It was assumed that through learning the weights within the assumed spatial priority map are up- and down regulated optimizing attentional selection. We have argued that in conditions in which one event predicts an upcoming event the weights within this spatial priority map need to be dynamically (from trial to trial) adjusted (Li et al., 2022; Li & Theeuwes, 2020). The current study provides direct evidence for these dynamic weight changes within the priority map as the presentation of a particular stimulus generated neural patterns at the location where the next stimulus was expected to be presented. Participants learned an arbitrary spatial sequence consisting of 4 dots (A-B-C-D), and after learning, partial sequence trials, wherein only a single dot (e.g., - B - -) appeared at its expected moment in time, were intersected with full sequence trials. Using the posterior electrodes, we were able to decode the spatial position of expected, yet critically omitted stimuli within these partial sequences, visualizing the dynamic weight changes within the presumed spatial priority map on a trial-to-trial basis. Our findings are fully consistent with those of Ekman et al. (2023) who used fMRI and showed reactivations of future sequence locations in both V1 and hippocampus.

The present findings contribute to an extensive body of literature demonstrating that learned associations are actively utilized to predict the imminent future (e.g., Dayan, 1993; Ekman et al., 2023; Fang et al., 2023; Stachenfeld et al., 2017). However, studies investigating how expectations induce anticipatory activity in early visual cortices (e.g., Ekman et al., 2017; Gavornik & Bear, 2014; Hindy et al., 2016; Kok et al., 2012) have yet to clarify whether such anticipatory traces represent expected future states in a temporally discounted fashion or solely reflect the learned association between different stimuli without the added complexity of a temporal structure (e.g., Deuker et al., 2016; Horner et al., 2015; Rolls, 2016). While Ekman et al.’s observation that events in the close future were more prominently represented than events in the more distant future supports the former account, in that study, the limitations of the BOLD approach made it impossible to provide a detailed analysis of how these learned predictions evolve over time. In our study, leveraging the high temporal resolution of EEG in conjunction with multivariate analysis, we demonstrate that any stimulus in a fixed, yet passively viewed sequence triggers the internal representation of the immediately following stimulus in the sequence at its expected moment in time. Crucially, this above-chance classification of an expected yet omitted stimulus was achieved by an independent pattern estimator trained on the evoked signal elicited by stimuli presented in random order, thereby avoiding confounding effects related to prediction. This confirms not only that future representations are represented in a temporally discounted fashion but also that the predicted future recruits similar neural processes as when viewing those stimuli.

The finding that learned spatial associations could evoke sensory-like stimulus representations in the absence of sensory input aligns with prior studies demonstrating decoding of anticipated positions during circular object motion (i.e., clockwise or counterclockwise) around fixation. However, there is a crucial difference between previous studies using rotating sequences and the current one. Critically, in previous studies, the rotating stimulus either induced illusory motion (Hogendoorn & Burkitt, 2018) or participants were explicitly instructed to mentally envisage the subsequent dot position each time a sound was played (Robinson et al., 2020). In contrast, our stimuli were not only presented in an arbitrary visual dot sequence, but this sequence also did not necessitate active planning to anticipate different future outcomes. Instead, the sequences were passively viewed, and therefore it is unlikely that the internal representations are the result of top-down attention; instead it is likely that they are formed automatically and incidentally via statistical learning (Christiansen, 2019; Frost et al., 2019). It is generally assumed that statistical learning operates through local weight changes in assumed spatial priority maps, enhancing the weights associated with locations likely to contain relevant information and downregulating those with a higher distractor probability (e.g., Fecteau & Munoz, 2006; Ferrante et al., 2018; Theeuwes et al., 2022; Zelinsky & Bisley, 2015). To date, studies examining local weight changes within the priority map have did so in paradigms where the learned locations, be it targets or distractors, were static across stretches of time (Duncan et al., 2023; Ferrante et al., 2023). By contrast, here the dot stimulus moved across space, necessitating a flexible priority landscape. At a behavioral level, we have previously shown that while learned attentional biases can be persistent, the priority landscape can also be flexibly adjusted across trials if needed, enhancing efficiency when a target at a specific location is consistently preceded by a target at the opposite display location (Li et al., 2022; Li & Theeuwes, 2020). Although the current study did not require attentional selection for each individual event, attention systematically moved from one location to the next, over time, activating the expected event in response to the current event.

Although we observed robust generalization from the random pattern estimator to both present as well as omitted dots within the structured sequence, there were clear differences in the temporal classification dynamics. Whereas decoding of viewed dots was largely restricted to the diagonal, suggestive of a chain of distinct generators (King & Dehaene, 2014), generalization to internally generated representations mostly relied on two training intervals (Figures 4c and 5a): one starting at around 100 ms and the other starting at approximately 200 ms. In the context of spatial attention, this early interval is related to early sensory/perceptual processing (indexed by P1 and N1 ERP components, e.g., Hillyard & Anllo-Vento, 1998), whereas the second interval correlates with post-perceptual processes such as attentional selection (indexed by N2pc ERPs component, e.g., Luck & Hillyard, 1994). It thus appears that both sensory/perceptual and post-perceptual processes underpin the activation of internal representations for the anticipated yet omitted stimuli. The fact that these processes generalized to relatively large stretches in time in stimulus absent conditions is not surprising given that in contrast to evoked signals, internally generated processes are likely to be much more variable, resulting in more diffuse time-locked signals (see also Robinson et al., 2020). In line with this explanation, the decoding accuracy of present dots was consistently higher when the dot was embedded within a full sequence compared to the same dots presented in isolation. This effect coincided with a reduction in the N1 elicited by dots in partial sequences. It suggests that attention could be precisely tuned to each individual event in full sequences, while this attentional tuning became more variable when dots were omitted from the sequence, and participants had to rely on internally generated representations. The diffuse nature of these internally generated representations might also explain why, in contrast to the Ekman et al. (2023), we could only decode the immediate successor representation, but not the subsequent representations within the sequence. Although these representations may have been present in the signal, the jitter caused by increasing temporal uncertainty arguably attenuated the signal to a degree that prevented their retrieval by our pattern estimator.

It is generally assumed that the neural signals elicited by successive stimuli with longer ISIs are less contaminated than those with shorter ISIs. However, to date this assumption has not been empirically tested. In a recent study by Grootswagers et al. (2019), hundreds of different images were presented at two different rates without an ISI (i.e., 20 Hz and 5 Hz). It was observed that decoding performance was higher and lasted longer in the 5-Hz condition compared to the 20-Hz condition, with no difference in the onset time between the two. Notably, the decoders captured category features rather than low-level features, as determined by the cross-validation procedure that should generalize to the test images. In comparison, the present study focused on decoding a specific low-level feature (i.e., position), making a comparison between two relatively slower frequencies. Consequently, our findings align with those of Grootswagers et al. (2019), and also provide some suggestions regarding ISIs for future decoding studies, especially those aim at decoding low-level features like positions and orientations. While the benefit of short-ISI pattern estimator sequences is that they are less time consuming, if time permits it may be best practice to use a pattern estimator with longer ISI’s as these provide a better model of stimulus specific representations.

In conclusion, the present study successfully decoded the positions of omitted anticipated dots within a learned spatial sequence, visualizing the dynamic weight changes within the presumed spatial priority map on a trial-to-trial basis. We demonstrated that the internal spatial representations, shaped by prior experiences, involved both sensory/perceptual and post-perceptual processes. Our research highlights the brain’s remarkable ability to dynamically prioritize and internally generate representations for anticipated locations in a future-oriented manner, even in the absence of sensory inputs. These findings help us further understand how the brain predicts and optimizes responses to external stimuli in a constantly changing world.

## Methods

### Participants

The sample size was determined based on previous EEG studies decoding dot positions in a circular configuration (Blom et al., 2020; Hogendoorn & Burkitt, 2018; Robinson et al., 2020). Twenty-five undergraduate students (22 females and 3 males; *M*age = 21.64 years, *SD*age = 2.36) recruited from research pool at the Vrije Universiteit Amsterdam participated in the study in exchange for course credit or monetary compensation. Two participants were replaced: one due to a poor connection to the earlobe channels, and the other one because of a recording error. All participants reported normal or corrected-to-normal visual acuity and gave informed consent prior to the start of the experiment. The study was approved by the Ethical Review Committee of the Faculty of Behavioral and Movement Sciences of the Vrije Universiteit Amsterdam.

### Apparatus

The experiment was created on a Dell Precision 3640 Windows 10 computer equipped with an NVIDIA Quadro P620 graphics card in OpenSesame (Mathôt et al., 2012) using PsychoPy (Peirce, 2007) functionality. Stimuli were presented on a 23.8-inch LED monitor (ASUS ROG Strix XG248Q; resolution: 1920 × 1080 pixels) at a refresh rate of 240 Hz. Participants were seated in a dimly lit room with their head positioned on a chin rest at a viewing distance of approximately 65 cm. Behavioral responses were collected via a standard keyboard. EEG signals were recorded using ActiView software (Biosemi, Amsterdam, Netherlands), and as common for Biosemi, two additional electrodes were used as reference and ground electrodes during recording. An Eyelink 1000 eye-tracker (SR Research, Ontario, Canada) sampling at 1000 Hz was used to on- and off-line monitor the gaze positions of both eyes during the experiment. Participants heard a low-volume beep sound from the speaker whenever their fixation deviation exceeded 1.5 degrees of visual angle (dva). Each time the participant moved away from the chinrest, the eye-tracker was recalibrated via a five-dot calibration procedure until the spatial error for each dot position was below 1 dva. Drift correction was applied before each block started.

### Stimuli and Design

The paradigm was modeled after Ekman et al. (2023). As illustrated in Figure 1, all stimuli were presented on a grey (RGB: 128/128/128) background. To elicit reliable and stable fixation, a combination of bull’s eye and hair cross (Thaler et al., 2013) was used as the central fixation marker (diameter: 0.7 dva; RGB: 0/0/0) which remained visible throughout the entire experiment. The stimulus was a white circular dot (diameter: 1.2 dva; RGB: 210/210/210), which could appear at any of eight possible locations (i.e., 22.5°, 67.5°, 112.5°, 157.5°, 202.5°, 247.5°, 292.5°, 337.5° of polar angle from the vertical line, see Figure 1b) at 6.5 dva away from fixation. This dot, when present, remained on screen for 100 ms.

Individual dots were presented in sequence in different block conditions, the *main task* and *pattern estimator* blocks. In the main task, participants viewed a sequence of four successively presented dots (A-B-C-D). As illustrated in Figure 1a (top row), following Ekman et al. (2023), the sequence trajectory was created such that each quadrant was stimulated once per sequence and neighboring locations were thus never part of the same sequence. Specifically, the starting position of the first dot determined the sequence trajectory such that the second dot always appeared 180° away from the starting position; the third dot was presented 90° clockwise from the second position; the last dot appeared opposite from the third position. Each participant was assigned to one of eight possible sequences. Each dot was shown for 100 ms with an inter-stimulus interval (ISI) of 200 ms, and individual sequences were separated by a jittered inter-trial interval (ITI) between 500 and 750 ms. On a small subset of these sequences (i.e., 10% of full sequence trials, the last dot was rendered in grey (RGB: 150/150/150) rather than white and participants were instructed to detect these grey dot presentations by means of a speeded space bar press (timeout 700 ms), while holding fixation at all times. To ensure that these rare target trials were relatively evenly distributed across trials within each block, trial order was pseudo-randomized with the constraint that consecutive grey dots were separated by minimally five sequences. Each time a grey dot was not reported, the text “Be focused!” appeared in red for 500 ms at the center of the screen. Also, the ITI following all target present trials was extended by 300 ms to reduce the influence on the next trial. The sole purpose of this detection task was to keep participants engaged with the stimulus sequences and hence behavioral data were not analyzed, and epochs time-locked to grey dot onset were excluded from further analysis.

The main task was separated into four learning blocks and 12 main task blocks. Within each learning block, the full sequence was repeated 130 times (out of which 13 contained a grey dot at the final position). By contrast, in main task blocks, 88 full sequence trials (out of which 9 contained a grey dot at the final position), were intermixed with 44 partial sequences. These partial sequences were identical to full sequences, with the difference that only one of the four dots in the sequence (with equal probability) was actually displayed (see bottom of Figure 1a). The purpose of these partial sequences was to probe neural activations of expected events. Sequence order within these main blocks was pseudo-randomized with the extra constraint that a partial sequence trial must be preceded by a non-target full sequence and not followed by another partial sequence. Feedback containing the hit rate and average response times was given at the end of each block.

In addition to the main task blocks, we also ran separate pattern estimator blocks, which served as an independent source of training data for the multivariate decoding analyses. Within these blocks, successive sequences of dots at all 8 possible locations were shown in random order (see Figure 1c), with the restriction that a dot position was never repeated at the crossing between sequences. Participants were instructed to keep fixation, while passively viewing these randomly concatenated sequences.

EEG decoding based on independent pattern estimator blocks has been successfully applied, where the ISI in between successive stimuli was either short (range: 0-200 ms; e.g., Alilovic et al., 2021; Blom et al., 2020; Robinson et al., 2020) or long (range: 433-1200 ms; e.g., Hogendoorn & Burkitt, 2018; Kok et al., 2017), with each approach having their own benefits. It has been demonstrated that even a simple flashing stimulus, as also used here, can evoke cortical response for ∼1 s (Carlson et al., 2013; King & Wyart, 2021) and the advantage of pattern estimator with longer ISI’s is that the signal evoked by each individual stimulus is less contaminated by evoked responses from preceding stimuli. By contrast, the benefit of short-ISI pattern estimator, apart from being less time consuming, the high rate of stimulus presentations also makes them less susceptible to systematic eye movements. Since this is the first time, we have attempted to decode the spatial position of expected dots in the absence of visual input, which is arguably more challenging due to the event of interest not being time-locked to an evoked response, we chose to employ both type of pattern estimators to establish whether one type is more suited than the other. To balance the length of ISI on long-ISI blocks and the length of the entire experiment, we chose an ISI of 600 ms for long-ISI blocks and an ISI of 100 ms for short-ISI blocks.

The three long-ISI pattern estimator blocks (50 trials each, ∼4.7 min) were introduced at the start of the experiment such that the longer ISI’s were not contaminated by learned anticipations in response to structured sequences. Subsequently, participants completed four learning blocks (∼4 min/block) to introduce them to the structured sequence. Then, following previous work (Kok et al., 2017; Robinson et al., 2020) every short-ISI pattern estimator block (25 trials each, ∼0.7 min) was interleaved with two main blocks (i.e., intermixed full and partial sequences). In total there were thus 300 pattern estimator sequences (i.e., 8 unique positions in random order), divided into three long-ISI blocks and six short-ISI blocks. For each participant, the entire experiment including preparation lasted about three hours.

### Analysis scripts

All preprocessing and subsequent analyses were conducted in a Python environment, using custom-written scripts based on functions from MNE-python (Gramfort et al., 2013), PyGazeAnalyser (Dalmaijer et al., 2014) and scikit-learn (Pedregosa et al., 2011) modules. The scripts are available at https://github.com/dvanmoorselaar/DvM.

### EEG recording and preprocessing

EEG data were acquired at a sampling rate of 512 Hz, using a 64-electrode cap with electrodes arranged according to the international 10-10 system (Biosemi ActiveTwo system; and two external electrodes placed on earlobes (used as an offline reference). Horizontal and vertical electrooculogram (HEOG and VEOG) were recorded via external electrodes located ∼1 cm lateral to the outer canthi, and ∼2 cm above and below the right eye, respectively. Recordings were filtered online with a high-pass filter of 0.16 Hz and a low-pass filter of 100 Hz. As an initial preprocessing step, EEG data was referenced to the average signal recorded over the left and right earlobe, and subsequently a zero-phase shift finite impulse response (FIR) high-pass filter at 0.01 Hz (van Driel et al., 2021) was applied to remove slow drifts.

Before cleaning, malfunctioning electrodes (*M* = 0.6, *SD* = 0.91) marked during recording were temporarily removed. Continuous EEG signals was epoched from −100 ms relative to the onset of a full sequence until the end of that sequence. Given that we had sequences of various lengths this resulted in epochs of various lengths (−100 to 1200 ms for full and partial sequences; −100 to 1700 ms for short-ISI pattern estimator sequences; −100 to 5200 ms for long-ISI pattern estimator sequences), which were all preprocessed independently. All epochs were extended by 500 ms at the start and end of the epoch to control for filter artifacts during preprocessing. First, independent component analysis (ICA) as implemented in MNE (method=‘picard’) was fitted on 1-Hz high-pass filtered epochs to remove eye-blink components from the 0.01-Hz filtered epochs. Second, an automatic artifact-rejection procedure was used to detect electromyography (EMG) noise. Specifically, the epochs were band-pass filtered at 110−140 Hz and transformed into a z-score threshold per participant based on the within-subject variance of z-scores (de Vries et al., 2017). Epochs that exceeded the threshold within the time window of interest were flagged (Duncan et al., 2023). To decrease the number of false alarms, for each marked epoch, the five electrodes that contributed most to accumulated z-score were identified. Then these electrodes were interpolated one by one using spherical splines (Perrin et al., 1989), checking after each interpolation whether the data still exceeded the threshold. Epochs were only excluded if the z-score threshold was still exceeded after this iterative procedure, which resulted in an average rejection of 5.85% (*SD*: 6.44%) of full/partial sequences, 14.37% (*SD*: 11.25%) of long-ISI pattern estimator sequences, and 9.95% (*SD*: 12.39%) of short-ISI pattern estimator sequences. Then, temporarily removed malfunctioning electrodes were interpolated using spherical splines. As a final step, in all three individual datasets the epoched sequences were split into individual dot-based epochs by creating smaller epochs (i.e., −100 to 300 ms) relative to the onset of each individual dot, irrespective of whether that dot was present or absent.

To eliminate the influence of eye movements on EEG decoding, after preprocessing we also rejected dot-based epochs on the basis of deviations from fixation as detected by the eye-tracker. To this end, gaze position (i.e., x and y coordinates) were epoched relative to the onset of the sequence using the same windows used during EEG preprocessing. Missing data due to blinks detected by the eye-tracker were interpolated by padding 100 ms before and after the blink interval (according to table 2 in Hershman et al., 2018). To control for drifts in the eye-tracker data, eye epochs without a saccade in the 100-ms pre-trial window were shifted toward fixation. The epoched data was then down-sampled to 512 Hz and aligned to the corresponding EEG epochs by creating dot-based epochs. Each of these individual dot-based epochs were summarized by a single value indicating the maximum fixation deviation measured in continuous 40-ms intervals of data (van Moorselaar & Slagter, 2019). If this value was above 1 dva, this epoch was marked for exclusion. In case of missing eye-tracker data, we used a sliding-window step method (length: 200 ms, step: 10 ms, threshold: 20 μV) on HEOG to detect eye movements. Combining these two methods, we excluded an average of 5.51% (*SD*: 5.1%) of sequence epochs, 4.59% (*SD*: 4.47%) of long-ISI pattern estimator epochs, 3.46% (*SD*: 4.33%) of short-ISI pattern estimator epochs.

### Decoding analyses

To investigate the position representation of both present and absent dots on the screen, we applied multivariate pattern analysis, using linear discriminant analysis (LDA) in combination with a 10-fold cross-validation scheme, with channel voltages from the 29 posterior electrodes (i.e., CPz, CP1, CP2, CP3, CP4, CP5, CP6, TP7, TP8, Pz, P1, P2, P3, P4, P5, P6, P7, P8, P9, P10, POz, PO3, PO4, PO7, PO8, Oz, O1, O2, Iz) used as features (e.g., Chota et al., 2023; Hajonides et al., 2021; Robinson et al., 2020; Wolff et al., 2017) and the four dot positions of the full sequences as classes. Before classification EEG data was averaged across three epochs (i.e., matching dot positions), while ensuring that the number of observations per stimulus position that entered the analysis was balanced across classes so that training was not biased towards a specific class. Event averaged data was then baseline corrected using a −100 to 0 ms window and down-sampled to 128 Hz (to decrease the computational time of decoding analyses). To further increase signal-to-noise ratio (Grootswagers et al., 2017), the data were transformed per sample using principal component analysis (PCA) with 99% of the variance. Before applying PCA, the data was standardized. Note that while performing standardization and PCA on both the training and test data, the transformation statistics were estimated using the training data only. Next, within-block decoding was performed via a 10-fold cross validation procedure, such that at each time point, the classifier was trained on nine folds and tested on the remaining fold until each fold was tested once. Classifier performance was then averaged over folds and characterized via Area Under the Curve (AUC), where a value of 0.5 is considered chance classification.

These initial decoding analyses, which were performed separately for short- and long-ISI pattern estimators, served to establish whether the pattern estimator could reliably track the dot position on screen. To subsequently assess whether absent dots were represented, if at all, differently than present dots, in a next set of analysis, the classifiers, with the same parameters used during within-block pattern estimator decoding, were trained on all events from the pattern estimator sequences and tested on both present and absent dots in full and partial sequences. Considering that there was no dot on screen in the absent condition, in these analyses we adopted a temporal generalization approach, training and testing was performed at all combinations of time points, to assess whether and when the EEG signal contained information regarding anticipated, yet absent events (King & Dehaene, 2014). In doing so, we again ensured that the number of observations in both the training and test set were balanced per stimulus position to avoid inflated above-chance performance resulting from a bias in the distribution of classes (Fahrenfort et al., 2018; van Moorselaar et al., 2020).

### Event-related potentials (ERPs)

To isolate stimulus-evoked ERPs components, separate contralateral waveforms were computed for full and partial sequence conditions, using a set of 8 posterior electrodes (i.e., P1, P3, P5, P7, P9, PO3, PO7, O1 or P2, P4, P6, P8, P10, PO4, PO8, O2) (e.g., Hajonides et al.,2021; Luck et al., 1990; Luck & Hillyard, 1994). In doing so, the first dot position (i.e., A) was excluded as at that point in the sequence there is no difference yet between full and partial sequences. Epochs were baseline corrected using a 100 ms window preceding dot onset. Resulting waveforms were averaged across dot positions (i.e., BCD).

### Statistics

Decoding scores were evaluated across time via cluster-based permutation tests with one sampled two-sided t-tests with cluster correction (*p* = .05, 1024 iterations; chance level = 0.5) using MNE functionality (Gramfort et al., 2013). The same procedure, but with a paired sampled t-test, was used to compare the difference between short- and long-ISI pattern estimators across time with the time window of the long-ISI pattern estimator matched to the short-ISI pattern estimator window.

## Author contributions

Ai-Su Li, Conceptualization, Methodology, Validation, Formal analysis, Investigation, Resources, Data curation, Funding acquisition, Visualization, Project administration, Writing - original draft, Writing - review and editing; Jan Theeuwes, Conceptualization, Funding acquisition, Writing - original draft, Writing - review and editing; Dirk van Moorselaar, Conceptualization, Methodology, Software, Validation, Formal analysis, Investigation, Resources, Visualization, Supervision, Writing - original draft, Writing - review and editing.

## Open practices statement

Data will be available at https://github.com/AisuLi/dot_seq, and the experiment was not preregistered.

## Acknowledgements

AL was supported by National Key Research and Development Program of China (2022YFB4500601) and National Natural Science Foundation of China (32171049). DvM and JT were supported by a European Research Council (ERC) advanced grant 833029 –[LEARNATTEND].

## Supplementary materials

**Figure 2—figure supplement 1.**
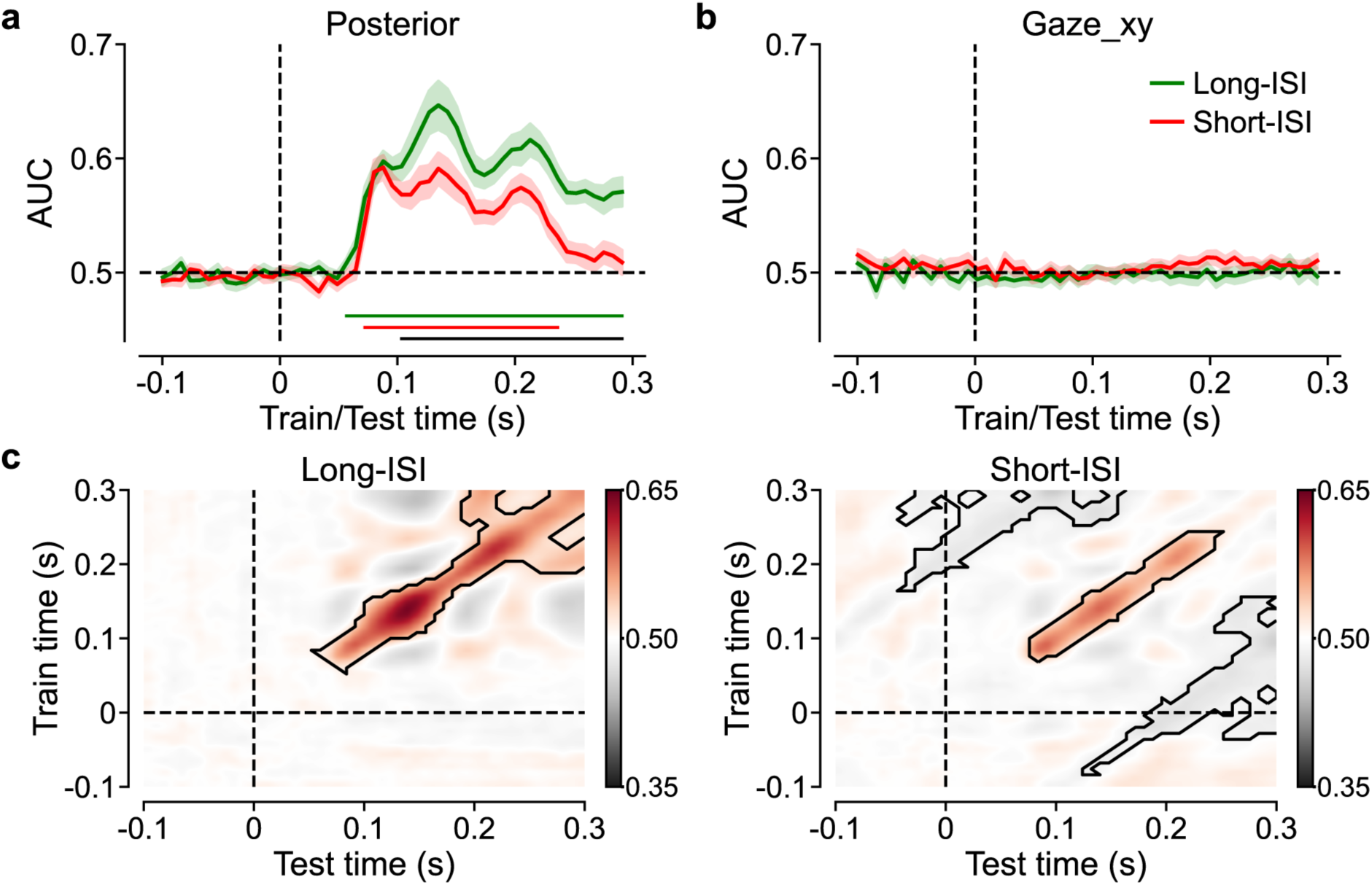
Pattern estimator position (all eight possible positions) decoding results. The decoder was trained and tested at the same time point with posterior electrodes (a), or gaze positions (b) as features. Horizontal dashed lines indicate chance level while vertical dashed lines indicate dot onset. Green and red solid lines represent long- and short-ISI pattern estimators, respectively. Shaded region represents standard error of the mean (SEM). Corresponding color bars below the x-axis indicate clusters that significantly differed from chance while the black bar indicates clusters with a significant difference between long- and short-ISI pattern estimators (*p* < .05). (c) Temporal generalization matrix, with posterior electrodes as features, showing classification performance as a function of training and test time point in long- and short-ISI blocks. Solid black lines indicate clusters that significantly different from chance (*p* < 0.05).

## Notes

### Competing Interest Statement

The authors have declared no competing interest.

